# Vegetation and Microbes Interact to Preserve Organic Matter in Wooded Peatlands

**DOI:** 10.1101/2020.04.11.034108

**Authors:** Hongjun Wang, Jianqing Tian, Huai Chen, Mengchi Ho, Rytas Vilgalys, Xingzhong Liu, Curtis J. Richardson

**Author notes:** These authors contributed equally to this work (Hongjun Wang, Jianqing Tian). **Correspondence and requests for materials** should be addressed to H.W. or C.J.R.

## Abstract

Peatlands have persisted as massive carbon sinks over millennia, even during past periods of climate change. The commonly accepted theory of abiotic controls (mainly anoxia and low temperature) over carbon decomposition cannot explain how vast low-latitude wooded peatlands consistently accrete peat under warm and seasonally unsaturated conditions. Similarly, that theory cannot accurately project the decomposition rate in boreal peatlands where warming and drought have decreased *Sphagnum* and increased shrub expansion. Here, by comparing composition and ecological traits of microbes between *Sphagnum*- and shrub-dominated peatlands, we present a previously unrecognized natural course that curbs carbon loss against climate change. Slow-growing microbes decisively dominate the studied wooded peatlands, concomitant with plant-induced, high recalcitrant carbon and phenolics. The slow-growing microbes inherently metabolize organic matter slowly. However, the fast-growing microbes that dominate our *Sphagnum* site (most boreal peatlands as well) decomposed labile carbon >30 times faster than the slow-growing microbes. We show that the high-phenolic shrub/tree induced shifts in microbial composition may compensate for positive effects of temperature and/or drought on metabolism over time in peatlands. This biotic self-sustaining process that modulates abiotic controls on carbon cycling may help better project long-term climate-carbon feedbacks in peatlands.

## Introduction

Peatlands cover only 3% of land surface but currently maintain 600–700 Gt of carbon (1 Gt = 10^15^ g), which exceeds global vegetation carbon stores and is close to the pool of atmospheric CO2 (1, 2). Hence, both the fate of the massive carbon stores in peat and the way peatlands, particularly their carbon-sequestration/release processes, respond to climate change are highly important to future climates. Generally, rates of carbon decomposition via soil microbial respiration increase exponentially with rising temperature in the short term (3). Climate warming and drought not only increase peat loss by accelerating decomposition but also cause substantial losses of the keystone mosses like *Sphagnum* in the vast boreal peatlands, followed by shrub expansion and its uncertain effects (4-6). Such cascading events could invoke a catastrophic positive feedback to global warming (7-9). However, long-term warming experiments in grasslands (10, 11) and studies spanning a wide range of mean annual temperature (MAP) globally (12, 13) show declining microbial metabolism over time under experimental warming or in warmer regions. By now most experimental studies in peatlands have lasted only for months to decades, and such time scales are deemed too short to detect long-term (>100 years) effects of climate change on millennial peatlands that may have complex evolutions/successions during past climatic fluctuations (14). High-resolution stratigraphic analyses on peat profiles across boreal areas documented that vegetation composition and net primary productivity played key roles in carbon accumulation during the last millennium (14, 15). We compiled soil respiration data from >150 peatland sites across latitudes between 2°S and 75°N (Dataset 1) to test whether the dependence of decomposition on temperature applies to a wide range of mean annual temperature (MAT) in peatlands. As both heterotrophic and autotrophic respirations were included here and plants with higher biomass in the tropical regions beget higher autotrophic respiration (16), we expected to see an apparent exponential rise in soil respiration along with increasing MAT. However, we found the idea unsustainable (Fig. 1). The paleontological evidence (14, 15) and our large-scale soil respiration analysis challenge the current abiotic-factor-dependent peat decay models (mainly temperature and water level) that is embedded into the Earth System Models to project climate-carbon feedback (7, 9). This disagreement, we assume, could result from the latent role of changing plant communities and their associated ecological and biogeochemical processes—a commonly occurring state shift in peatlands induced by persistent climate change (14). Changes in plant communities among mosses, sedges, shrubs and trees may bring forth substantial top–down and bottom–up regulations (17) on the peatland ecosystem through alteration of plant–microbe traits, specifically plant/soil chemistry (18) and microbial composition/function (5, 19-21). Although temperature dominantly controls microbial metabolism of soil carbon in monoculture or a constant environment, the long-term feedback between climate change and soil carbon processes (especially some evolutional acclimations in plant/microbial physiology and community composition in peatlands (11, 19-24)) is still unclear and possibly one of the major uncertainties and challenges in projecting future climate in the Earth System Models(25). Thus, recognition of the biotic control can be central to the development of a meaningful framework for unravelling the responses of the perpetual peatlands to climate change (19, 21).

**Fig. 1.**
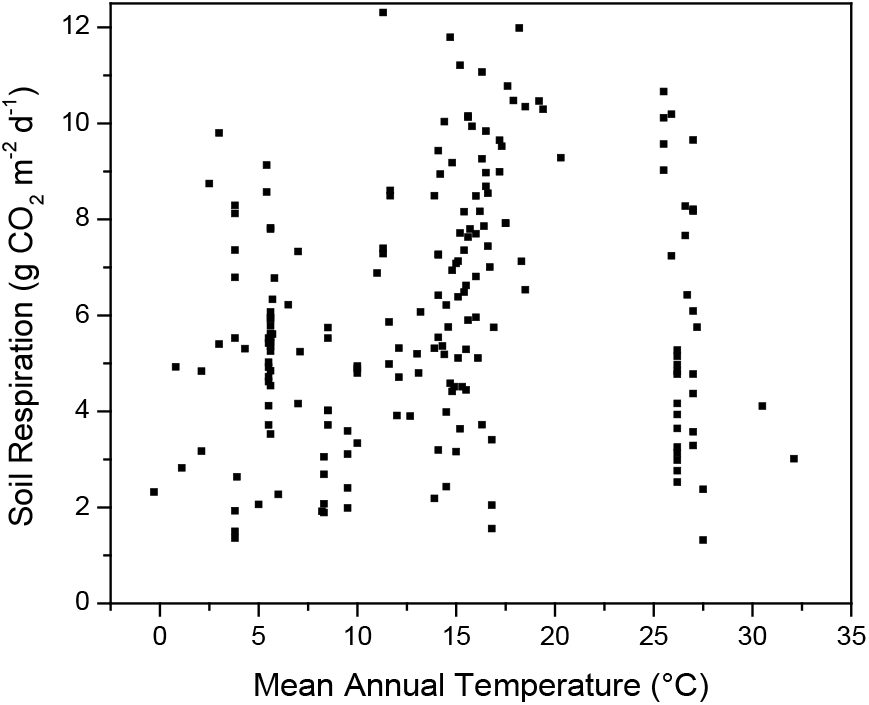
Mean soil respiration from boreal to tropical peatlands. Data were compiled from 31 published studies that contain annual mean soil respirations observed in >150 sites globally (Dataset 1).

## Results and Discussion

We used space-for-time substitution as an alternative to study shifts of ecological processes in peatlands over time. An ombrotrophic boreal *Sphagnum* peatland and a subtropical shrub peatland in the U.S.A were first selected. The *Sphagnum* site is located in the Marcell Experimental Forest, Minnesota (47°30’N, 93°27’W), USA, and the shrub site is in Pocosin Lakes National Wildlife Refuge, North Carolina (35°40’N, 76°32’W), USA. The *Sphagnum* site is dominated by *Sphagnum* mosses with scattered shrubs and black spruce (*Picea mariana*), while the shrub site has responded to climate change over the past 12,000 years through a transition of plant communities from boreal *Sphagnum*/spruce during late-glacial to the modern ericaceous shrubs today (6, 26). These shrub species in pocosins are similar to the expanding shrubs found in *Sphagnum* peatlands across boreal areas (4, 5). Our pilot results in the *Sphagnum* site are in line with the observations in many boreal peatlands, while such studies in low latitude system are still rare. Hence, two wooded peatlands were further tested in subtropical southern China and temperate western Canada.

To determine the underlying microbial communities and their ecological traits, we collected triplicate soil cores from the hollows and hummocks in the boreal *Sphagnum* peatlands and from 3 sites with different plants and water levels in the subtropical shrub peatlands (Table S1) and measured the composition and abundance of fungi—the predominant peat decomposers—and associated physiochemical peat parameters. Compared to the *Sphagnum* peat, dissolved phenolics in the shrub peat were about 6–10 times higher (Table S1). The *Sphagnum* and shrub peatlands distinctively differed in the fungal community composition (Fig. 2). At the *Sphagnum* site, the fungal groups were dominated by *Pseudeurotium*, Saccharomycetales and *Mortierella*, whereas at the shrub site, by *Archaeorhizomyces* and Helotiales (Fig. 2, Dataset 2). As microbial growth rates significantly affect carbon turnover in soil (27, 28), we roughly classified the dominant fungi as either fast- or slow-growing groups based on known growth traits of culturable species from each taxonomic group (Table S2). This is the best way we can approach by now although some inevitable biases exist in this classification. Notably, nearly 85% of fungi were likely fast-growing at the *Sphagnum* site, but at the shrub site a mere 2% were categorized as fast-growing and about 75% as slow–growing based on their ecological traits (Fig. 2, Table S2). Consistent with the majority of fungi found in many boreal peatlands (29), the dominant fungi at the *Sphagnum* site are members of pioneer saprobe communities that use simple carbon compounds and possibly possess *r*-selected strategy with fast growth rates(27-29). Unexpectedly, 11–97% of fungal sequences at the shrub site were assigned to *Archaeorhizomyces*, but only 0.4% on average at the *Sphagnum* site. *Archaeorhizomyces*, which represents one of the ubiquitous lineages of soil fungi, is also characterized by markedly slow growth (30). Another dominant fungal group (~5%) in the subtropical shrub site is the Helotiales, which includes certain fungi that form ericoid mycorrhizal fungi with resistant melanized cell walls (so called dark-septate endophytes) which are also characterized by slower growth rates (31). To determine what might control the fungal composition, we examined the relationship between fast-versus slow-growing fungal richness and physicochemical variables including dissolved organic carbon, dissolved phenolics, soil pH, soil moisture, NO_3_^−^–N and NH_4_^+^–N. Dissolved phenolics likely acted as the overarching regulator, not only directly limiting microbial activities (6, 32) but also allowing slow-growing fungi to thrive while hampering fast-growing fungi (Fig. 3). We also found soil acidity positively related to the relative richness of slow-growing fungi while negatively related to that of fast-growing fungi (Fig. S1). As there was a significant negative relationship between soil pH and dissolved phenolics (*r*^2^ = 0.428, *P*<0.0001), the dissolved phenolics in these peatlands might be mainly phenolic acids that increase soil acidity, thus further worsening extreme conditions that benefit the slow-growing fungi(28) or under which only slow-growing fungi can survive.

**Fig. 2.**
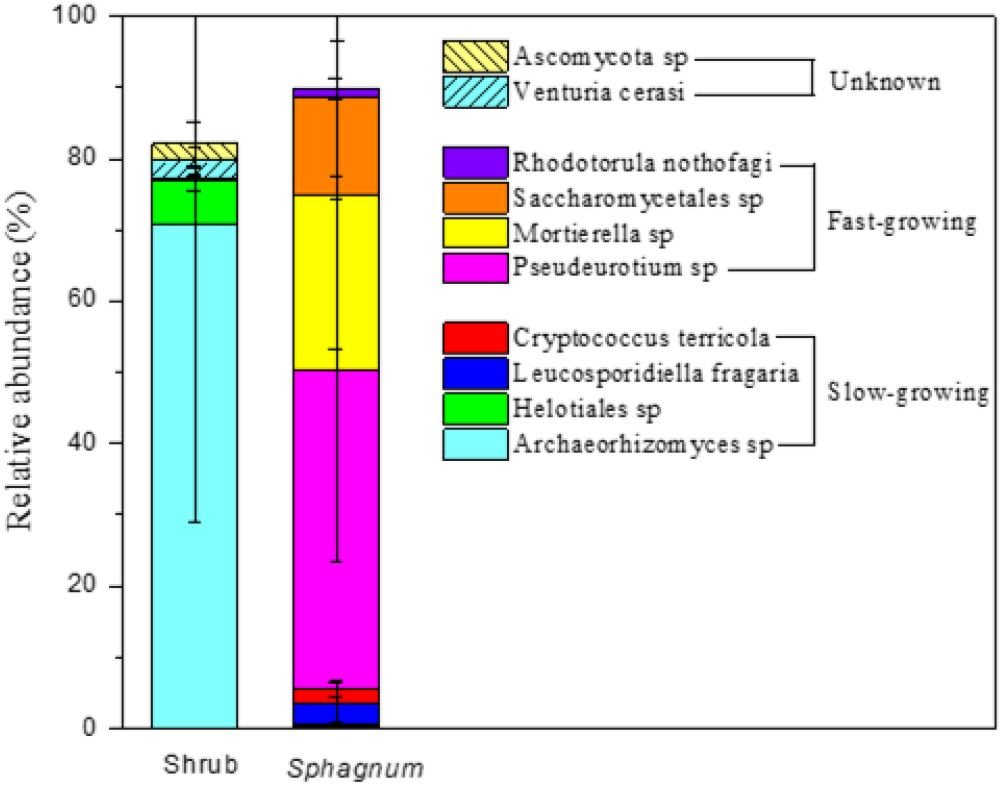
Dominant fungal composition (relative abundance >1%, see Dataset 2 for detailed OTUs distribution) and relative abundance (mean ± S.E.) of slow-growing and fast-growing fungi (see Table S2 for ecological traits of the dominant fungi) in the subtropical shrub and the boreal *Sphagnum* peatlands.

**Fig. 3.**
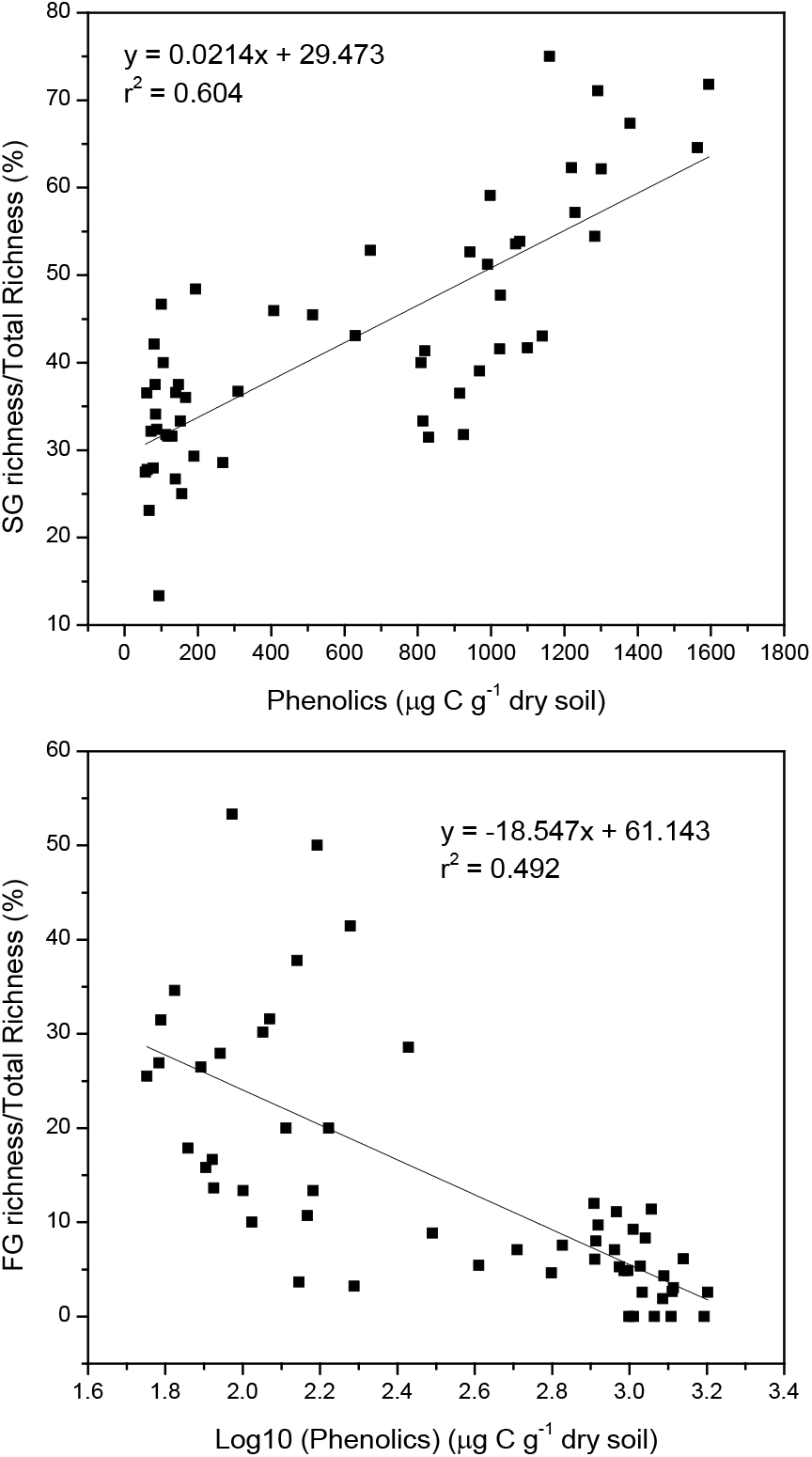
Effects of phenolics on ratios of slow- and fast-growing fungal richness to total richness. Species richness is the number of total OTUs observed in each sample. SG = slow-growing fungi, FG = fast-growing fungi.

Although we did not measure bacterial composition in this experiment, recent measurements (33, 34) at several of the same shrub and *Sphagnum* sites show that oligotrophic slow-growing Acidobacteria (27) dominated both peatlands. The relative abundance of Acidobacteria in the shrub sites (57%) was substantially higher than that in the *Sphagnum* sites (36%). Moreover, the fast-growing bacteria—including Betaproteobacteria and Bacteroidetes as copiotrophs (27) — were nearly absent from the shrub site (33) but contributed about 9% and 4%, respectively, in the *Sphagnum* site (34).

It is impossible to measure the relative growth rates of the dominant microbes in this study, because so many of the fungi are unculturable. To further test and verify the consequences of these likely fast-growing versus slow-growing microbes, we compared soil respiration rates per gram microbial biomass carbon (MBC) (12, 24) in the *Sphagnum* and the shrub peats through a reciprocal inoculation experiment using inocula and peat materials from both sites. We also added each inoculum to labile carbon-enriched mineral soil to test how the microbes decompose labile carbon. All soil media and incubators were sterilized by an autoclave before inoculation. Consistent with the microbial growth traits found in the literatures (Table S2), the microbes from the shrub peat decomposed both labile and complex organic material at much slower rates than the microbes from the *Sphagnum* peat (Fig. 4). Although the shrub peat is more highly recalcitrant than the *Sphagnum* peat, the fast-growing microbes from the *Sphagnum* inoculum decomposed the shrub peat 4 times faster than the microbes from the shrub peat did, which means the decomposers ultimately determine the decomposition rate. Moreover, the soil respiration rates from *Sphagnum* inoculum were dependent on the source of carbon, with rates at 320, 838 and 2315 mg C hr^-1^ g^-1^ MBC in the shrub peat, the *Sphagnum* peat and the labile carbon-enriched mineral soil, respectively. Hence, the dominant decomposers in the *Sphagnum* peatlands are likely fast-growing copiotrophs that generally adapt to using available resource rapidly (28). In contrast, soil media—including labile carbon—did not impact the microbial activities from the shrub inoculum (71–92 mg C hr^-1^ g^-1^ MBC). These results support our hypothesis that microbial metabolism in high-phenolic shrub peatlands is slower and that such a growth trait of a specific microbial community might be inherent (28).

**Fig. 4.**
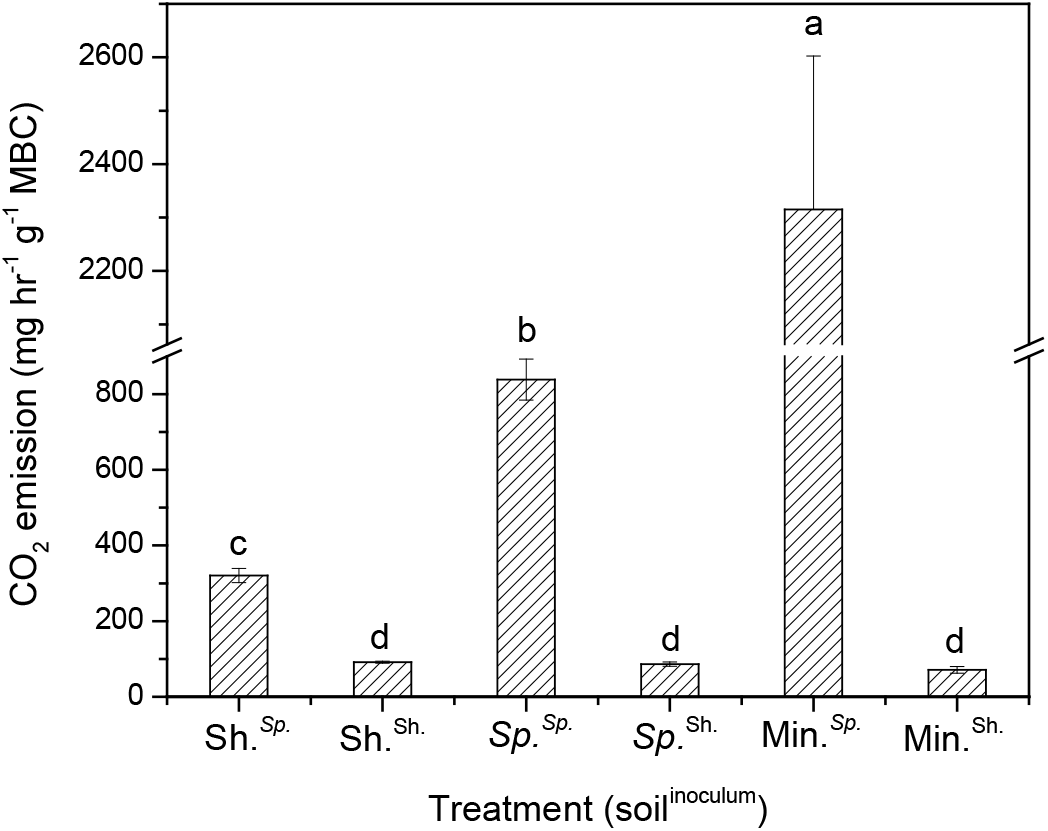
Standardized CO2 emission (mean ± S.E.) from sterilized soil media that include the boreal *Sphagnum* peat from Minnesota (*Sp*.), the subtropical shrub peat from North Carolina (Sh.) and the labile carbon–enriched mineral soil (Min.). All media were inoculated by inoculum made from the *Sphagnum*(^*Sp*^) or shrub (^Sh.^) peat. MBC represents microbial biomass carbon. Standardized CO2 emission was calculated based on the amount of MBC added to the soil media. Different letters indicate significant differences among treatments.

The slow-growing microbes that dominate the high-phenolic shrub site behave like *K*-selected taxa that may have outcompeted fast-growing *r*-selected taxa under steadily warmer and dryer conditions. The established slow-growing fungi, as well as bacteria (33, 34) lead to a lower carbon turnover in soil (6, 27). The dominance of the slow-growing microbes may explain why plant necromass does not completely decompose, but continues to accumulate as peat in low-latitude wooded peatlands, despite constant warming and frequent drought over millennia (6). This also further uncovers the observed slow decomposition under drought in the subtropical shrub peatlands (6), which was likely caused not only by the anti-microbe role of increased phenolics (6, 35) but also the magnified slow-growing decomposers induced by higher phenolics. Collectively, our field and lab experiments demonstrate for the first time that a phenolics-linked plant-microbe interaction acts as a natural curb on carbon loss in low-latitude wooded peatlands and will likely also in future boreal peatlands with climate-induced shrub expansion. This biotic self-sustaining process driven by consistent increases in temperature and drought over time appears to be more important than direct abiotic controls in regulating long-term carbon–climate feedbacks in peatlands, which is critical for understanding and modeling how ongoing climate change affects peatlands across the globe.

We postulate that these slow-growing microbes have adapted to high-phenolic acidic conditions and become the dominant microbes with inherent slow metabolic processes, a major underlying feature of high-phenolic wooded peatlands developed under warm and relatively dry conditions. This hypothesis is further supported by a temperature sensitivity experiment, a fungal composition measurement in a subtropical peatland in Asia and a reanalysis of fungal composition in a wooded peatland in Canada (36). Although higher microbial biomass carbon was present in the shrub peat (3.8 ± 0.2 mg C g^-1^ dry soil) in North Carolina relative to the *Sphagnum* peat (2.6 ± 0.3 mg C g^-1^ dry soil) in Minnesota, the decomposition rate of the shrub peat was much slower at the same temperature and displayed lower temperature sensitivity than the *Sphagnum* peat (Fig. S2). As we have shown the fast-growing microbes decomposed highly recalcitrant peat more quickly than the slow-growing microbes did (Fig. 4), thus current fast-growing microbes in most boreal peatlands (29) could decompose stored carbon more quickly when temperature is higher, but the microbes in peat soil may gradually shift to slow-growing species with increasing phenolic contents through shrub expansion under climate change (5) or even by adding wood litter—a new appearing geoengineering in degraded peatlands(35). A recent study showed that the relative abundance of slow-growing fungi (Helotiales and *Archaeorhizomyces*) was over 80% in a *Sphagnum* peatlands where ericaceous shrubs dominated the wooded cover in Meadowlands, MN, USA (37), which shows that the on-going shrub expansion in some boreal peatlands have shifted in sync from fast-growing to slow-growing microbes. In addition, we examined a shrub-dominant subtropical peatland with *Sphagnum* layer underneath in Shennongjia, China (31°29’N, 109°59’E), where the majority of fungal taxa (80.4%) are also found slow-growing (Fig. 5), e.g. *Archaeorhizomyces* spp. (26.5%) and *Cryptococcus* sp. (34.0%). We also reanalyzed the fungal composition in a bog forest and a *Sphagnum*-shrub mixed peat bog in the Pacific coastal temperate rainforest in Canada (36). Slow-growing fungi also dominated these wooded peatlands (Fig. S3, Table S2). Importantly there was a significant negative relationship between soil respiration and richness of slow-growing fungi (Fig. 6), which indicates that the slow-growing fungi regulate the carbon turn-over rates in the wooded peatlands. Our research suggests that slow-growing microbes, which are less effective in decomposition under warmer climates, may dominate most wooded peatlands, particularly in low-latitude areas. The dominance of slower-growing microbes in these wooded peatlands, no exponential rise in soil respiration with increasing MAT from boreal to tropical regions (Fig. 1), and the vast amount of carbon stored in tropical wooded peatlands(2, 38) together support this deduction, even though our study sites included only three wooded peatlands in North America and Asia.

**Fig. 5.**
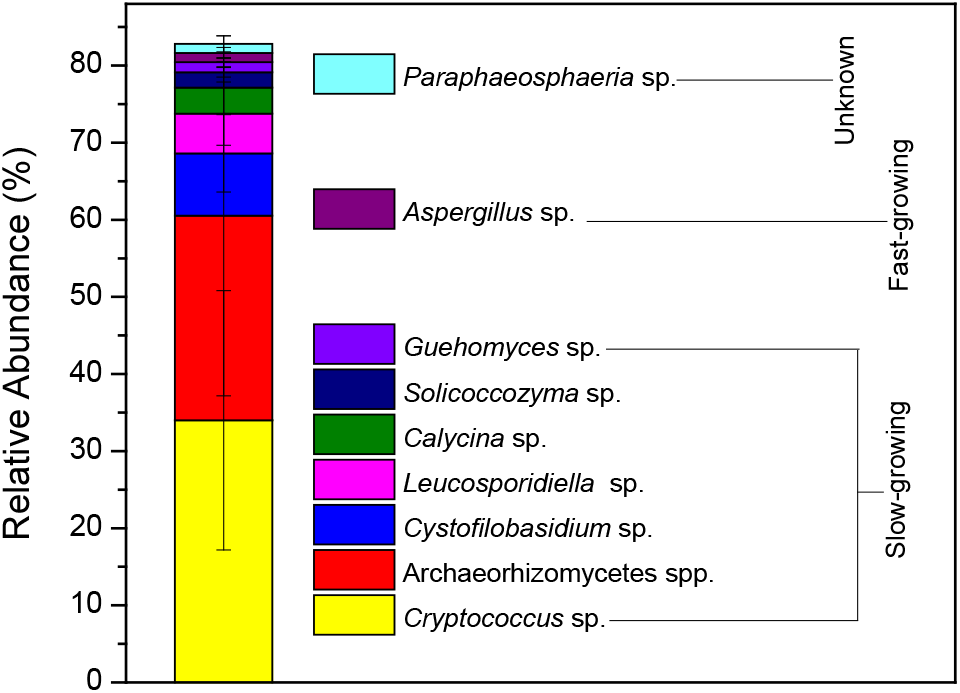
Relative abundance (mean ± S.E.) of the dominant fungi (OTU relative abundance >1%) in Dajiuhu subtropical peatland in China (see Table S2 for ecological traits of the dominant fungi).

**Fig. 6.**
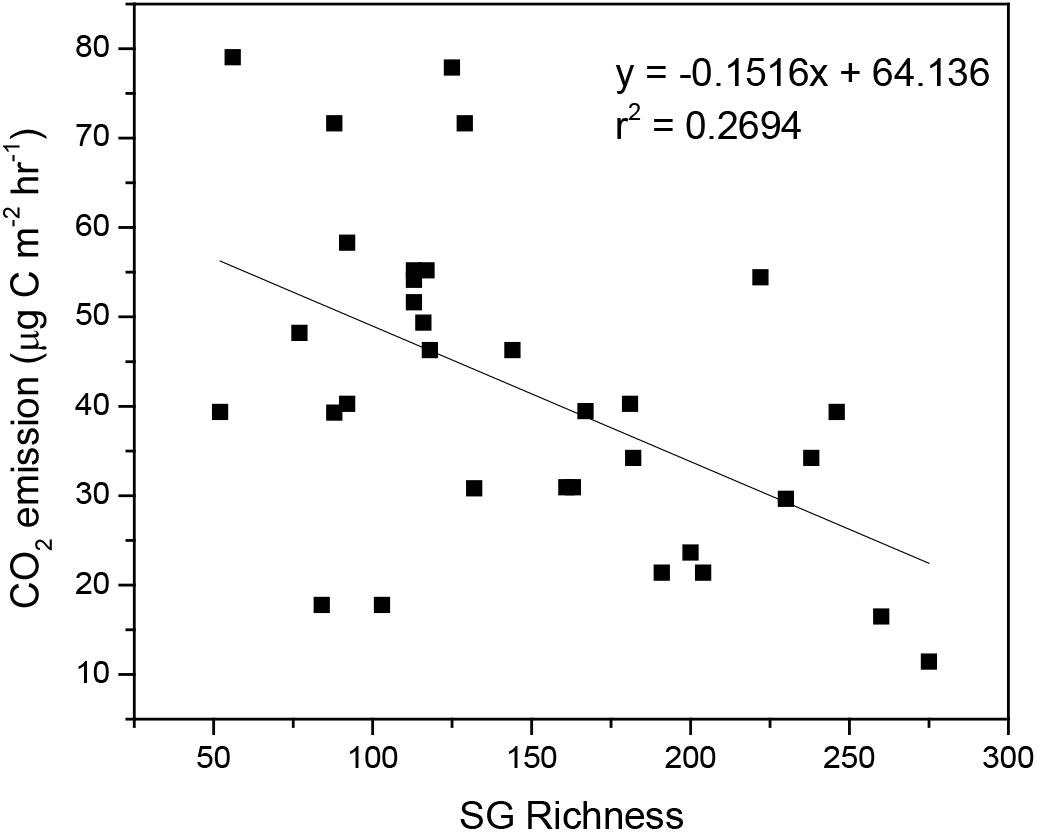
The relationship between richness of slow-growing fungi and CO2 emission in wooded peatlands in the Pacific coastal temperate rainforest in Canada. The richness of fungi was recalculated from raw amplicon reads in the European Nucleotide Arch Archive [ITS(ERS1798771-ERS1799064), CO2 emission data were collected from the Hakai Institute data repository at https://doi.org/10.21966/1.715630. SG = slow-growing fungi.

Finally, our findings provide new evidence that enduring peatlands that are highly resistant to increased temperature and drought may gradually shift to a new equilibrium state (19) with different microbes and plants that have adapted to the changed climate over time through their self-sustaining plant-microbe interactions (likely bridged by plan-induced phenolics). As biotic regulators, the co-shifting microbe and plant communities that were initially triggered by climate change appear to exert dominant controls on ecosystem C cycling and soil C sequestration, thus ensuring for continuing peat accretion in the new state. Our findings may have more immediate applications in carbon-climate feedback models and geoengineering strategies (35, 39). Embedding dynamic biotic factors into current abiotic-factor-dependent decay models could greatly advance the accuracy of the Earth System Models in projecting the fate of boreal peatlands with shifting plant/microbe communities (4, 5, 37) under climate change. This mechanism like a new framework would allow models to predict how the biotic processes of a peatland could modulate abiotic controls on the carbon cycle over time. Moreover, this mechanism further indicates that peatland geoengineering (39) adding high-phenolics natural materials like woody litter (35) could be an accelerated nature-based solution, similar to a natural state shift, preserving degraded peatlands not only in the short term through increasing phenolic contents (35) but also in the long term by encouraging phenolics-magnified, slow-growing microbes.

## Methods

### Study sites and soil sampling

Our major study sites were located in a shrub bog (6) in the Pocosin Lakes National Wildlife Refuge, North Carolina, USA and a *Sphagnum* bog (34) in the Marcell Experimental Forest in Minnesota, USA. Three sites (>1 km apart) around Lake Pungo including Pungo West, Pungo Southwest and Pungo East were selected at the shrub bogs. *Ilex glabra* and *Lyonia lucida* cover about 85% and 10%, respectively at Pungo West. *Ilex glabra* and *Lyonia lucida* are also dominate Pungo Southwest but distribute evenly, also there are many *Woodwardia virginica* during the growing season. The water level at Pungo Southwest is always higher than at Pungo West. Both Pungo West and Pungo Southwest have prescribed light fire every 4–5 years. There is no disturbance at Pungo East site over 30 years, where more dominant plant species exist, including *Lyonia lucida, Ilex glabra, Zenobia pulverulenta, Gaylussacia frondosa, Vaccinium formosum*. One hollows and one hummocks were selected at the *Sphagnum* bogs. A lot of mature trees including *Picea mariana, Pinus resinosa, Larix laricina* with different bryophytes and shrubs grow at both the hollows and the hummocks. *S. fallax* dominates the bryophyte layer at the hollows, and *S. angustifolium and S. magellanicum* dominate at the hummocks. The understory has a thick layer of ericaceous shrubs including *Ledum groenlandicum, Chamaedaphne calyculata, Vaccinium oxycoccos* at the hummocks, however, only scattered shrubs happen at the hollows. Other site information is described in Table S1. We took three soil cores at each sites (with a distance >4 m from each other), and each soil core was sliced to four sub-samples (0–5 cm, 5–10 cm, 10–15 cm, 15–20 cm). Big roots were removed in lab. The hair roots of all plants were included in the soil samples. Additionally, we took three soil cores at depth 0–10 cm in the shrub-dominant area in Dajiuhu peatlands in Shennongjia, China (31°29’N, 109°59’E) in May 2017. The dominant shrub at Dajiuhu is *Crataegus wilsonii with dense Sphagnum layer*. The samples were transported to the laboratory in iceboxes. Half of the samples were frozen at −80°C for DNA isolation; the other half was stored at 4°C for chemical analysis.

### Soil chemistry analysis

We used the deionized water extraction of fresh soil for dissolved organic carbon (DOC) and soluble phenolics measurements. DOC was measured as the difference between total C and inorganic C with a total C analyzer (Shimadzu 5000A, Kyoto, Japan). Soluble phenolics were measured by following the Folin-Ciocalteu procedure(40). Inorganic nitrogen (NH_4_^+^–N and NO_3_^−^–N + NO2^-^–N) extract with 2 M KCl was determined colorimetrically on a flow-injection analyzer (Lachat QuikChem 8000, Wisconsin, USA). Total carbon and nitrogen in soil were analyzed with combustion CN soil analyzer equipped with a TCD detector (ThermoQuest Flash EA1112, Milan, Italy). A 1:10 soil/water solution was used to measure soil pH.

### DNA extraction, PCR and sequencing

Genomic DNA was extracted from 0.25 g (fresh weight) of each homogenized soil sample using the PowerSoil DNA isolation kit (Mo Bio Laboratories, Carlsbad, CA, USA). DNA of each replicate was extracted three times and homogenized together as one DNA template. For Pocosin and Minnesota samples, a set of fungus-specific primers, ITS1F (3’- CTTGGTCATTTAGAGGAAGTAA-5’) and ITS4 (3’-TCCTCCGCTTATTGATATGC-5’), were used to amplify the internal transcribed spacer (ITS) region using barcoded ITS1F primers.

For Dajiuhu samples, ITS1F and ITS2 (3’- GCTGCGTTCTTCATCGATGC-5’) were used. All PCR reactions were repeated in triplicate, together with the negative controls in which the template DNA was replaced with sterile H_2_O. The amplicon concentration of each sample was determined after purification using Qubit^®^ 2.0 Fluorometer (Invitrogen, Grand Island, USA), samples pooled at equimolar concentrations, purified using AMPure Bead cleanup, and submitted to the core facility at Duke University (Durham, NC, USA) for sequencing using Illumina MiSeq (Illumina, San Diego, CA, USA).

### Bioinformatics processing

Sequence data of Pocosin and Minnesota samples are obtained from both ITS1 and ITS2 gene regions. ITS sequences were quality filtered and processed using the standard QIIME pipeline, with each fungal taxon represented by an operational taxonomic unit (OTU) at the 97% sequence similarity level. Singleton OTUs were omitted (41), and OTUs classified taxonomically using a QIIME-based wrapper of BLAST against the UNITE database(42, 43) (Supplementary Methods for further details). The quality and depth of coverage of both primers reads were not significantly different, thus libraries from ITS4 reads were used for further analysis of fungal communities. Taxonomic-based alpha diversity was calculated as the total number of phylotypes (richness) and Shannon’s diversity index (H’). A total of 150,967 ITS sequences from ITS2 region passed quality control criteria in the Pocosin and Minnesota sites. These sorted into 590 OTUs. Follow the same procedure, a total of 115,936 ITS1 sequences from Dajiuhu samples were assigned into 307 OTUs. Following the processing procedure described by Wilson(34), relative abundance of beta-proteobacteria at the controlled site in the boreal *Sphagnum* site was recalculated from Wilson and others’ sequence data(34) available from the National Center for Biotechnology Information at SRP071256. Relative abundance of fungi from a bog forest at the Calvert Island in Canada was recalculated from the raw amplicon reads in the European Nucleotide Archive, ITS (ERS1798771-ERS1799064).

### Lab incubations

We tested the decomposing capability of microbes in the *Sphagnum* and shrub peatlands by amending peat inocula from both sites in North Carolina and Minnesota to their peats and labile carbon-enriched mineral soil. Fresh *Sphagnum* and shrub peat inocula were prepared by mixing 0.5 kg of each type of fresh peat (10–20 cm) with 2 liters of deionized water. After 1 hour of stirring and 1-day settlement, the suspension liquid inoculum was filtered through a Buchner funnels (without filter, pore size 0.25–0.5 mm). We added 2 g of glucose to 50 g of nutrient-poor mineral soil (initially 0.05% total nitrogen, 0.64% total soil carbon) to produce a mineral soil medium with high labile carbon content. All incubation media (peat and mineral soil) and jars were sterilized by an autoclave before inoculation. About 30-g fresh *Sphagnum* peat (2.5–2.8 g in dry weight) or shrub peat (9.1–9.3 g in dry weight) or 50-g mineral soil with 2-g glucose was placed in Mason jars (triplicate, 8-cm diameter, 12-cm height, vacuum seal lid with a stainless steel fitting with sampling septum), then 20 ml of its own or other’s inoculum was added to the peat media, and 5 ml of inoculum from each site was added to the mineral soil. Finally, all samples were aerobically incubated at a constant temperature of 25°C. We initially used Parafilm M^®^ Laboratory film, which is air permeable but water resistant, to seal the top for 3-day equilibration, afterward we collected gas samples by syringe from the headspace of each jar at the beginning and end of 1-hour sealed incubation and used a GC (Varian 450, California, USA) to analyze CO2 concentration. As microbial biomass itself is a factor regulating soil respiration rates, standardized CO2 emissions at the microbial biomass were calculated based on the elevated CO2 concentration, time and air volume in the jar and the amount of added microbial biomass carbon from the inoculum. To prevent microbial acclimation to the assay chemistry (13, 24), we only incubated the soils shortly. A chloroform fumigation-extraction method (0.5 M K2SO4 to extract biomass C) (44) was used to determine soil microbial biomass carbon by the difference in measured carbon contents between fumigated and control replicates of each sample.

To test temperature sensitivity of soil respiration, nine fresh peat samples (30 g) from each site were added to jars and sealed with Parafilm M. Triplicate samples were incubated at 4°C, 25°C and 44.5°C. The highest temperature in this incubation does not match the *in-situ* conditions in our sites, but it may happen shortly in tropical wooded peatlands in future. After 3-day equilibration, we used the same method as above to measure gas emission and calculated soil respiration based on soil weight. We conducted regression analyses for soil from each site using R = *α*e^*β*T^, where R is soil respiration, coefficient *α* is the intercept of soil respiration when temperature is zero, coefficient *β* represents the temperature sensitivity of soil respiration, and T is soil temperature.

### Statistical analysis

One-way ANOVA with Duncan’s multiple-range test was used to compare the means. Standard error of the mean was calculated for each mean. The significant level of the test was set at a probability of 0.05. The ANOSIM function in the *vegan* package in R was used to test statistical significance in fungal composition within and among sites in the shrub peatlands and the *Sphagnum* peatlands (999 permutations), which shows that fungal communities were significantly different within sites at the shrub peatlands (Pungo East Pungo West and Pungo Southwest) and at the *Sphagnum* sites (hollows and hummocks) (Fig. S4).

### Data availability

The generated sequence data are available from the National Center for Biotechnology Information at SRP122579 and SRP158553. All other data in this study are available within the paper and its Datasets.

## Acknowledgements

We would like to thank Drs. David J. Levy-Booth and William W. Mohn at University of British Columbia and Dr. Joel E. Kostka at Georgia Institute of Technology for sharing their published data, Dr. William H. Schlesinger at the Cary Institute of Ecosystem Studies for his detailed comments on experimental design and data interpretation, Dr. Jeff Chanton at Florida State University, Dr. Dorothy Peteet at Columbia University, Dr. Christopher W. Schadt at Oak Ridge National Laboratory, Dr. Scott Neubauer at Virginia Commonwealth University, and Dr. Louis James Lamit at Syracuse University for their comments, Belen de la Barrera for laboratory measurement, Dr. Randy Neighbarger for technical editing. US DOE Office of Science, Terrestrial Ecosystem Sciences (DE-SC0012272), Key Research Program of Frontier Sciences, CAS (QYZDB-SSW-DQC007), the Duke University Wetland Center Endowment, and China Scholarship provided financial support.

## Author Contributions

H.W., J.T. and C.J.R. conceived the ideas and designed this research; C.J.R., H.W., R.V. and H.C. obtained funding; H.W., M.H., C.J.R., and H.C. collected field samples; J.T., H.C., and X.L. did microbial measurement and analyzed microbial community data; H.W. measured soil chemistry and conducted incubation experiments; H.W. and J.T. compiled data of soil respiration from literatures; H.W. wrote the manuscript with J.T. and C.J.R.; and all other authors discussed results and commented on the manuscript.

