## Supplementary methods, figures and tables for "Vegetation and Microbes Interact to Preserve Organic Matter in Wooded Peatlands"

(Supporting Information)

Hongjun Wang[1#*], Jianqing Tian[2,1#], Huai Chen[3], Mengchi Ho[1], Rytas Vilgalys[4], Xingzhong Liu[2], Curtis J. Richardson[1]

[1]Duke University Wetland Center, Nicholas School of the Environment, Box 90333, Duke University, Durham, NC 27708, USA.

[2]State Key Laboratory of Mycology, Institute of Microbiology, Chinese Academy of Sciences, Beijing 100101, China.

[3]Key Laboratory of Mountain Ecological Restoration and Bioresource Utilization and Ecological Restoration Biodiversity Conservation Key Laboratory of Sichuan Province, Chengdu Institute of Biology, Chinese Academy of Sciences, Chengdu 610041, China.

[4]Department of Biology, Box 90338, Duke University, Durham, NC 27708, USA.

[#]These authors contributed equally to this work.

**Methods**

**Figs. S1,S2, S3 and S4**

**Tables S1 and S2**

### Methods

#### Bioinformatic analysis

The trimming, sorting and quality filtering of reads were processed using the QIIME Pipeline(45). All sequences were demultiplexed into samples based on unique identification barcodes using the split_libraries_fastq.py script. As ITS1f and ITS4 are too long to pair-end, the sequences from both primers were processed separately. Sequences were sorted by gene region and primers were removed using cutadapt, and then reads were trimmed to 220 bp using -fastq_filter with a q10 minimum value as a quality filter. Sequences were dereplicated and singletons were removed. The ITS2 region from the remaining sequences was identified and extracted using the fungal ITSx software (46). Chimeras were detected and eliminated using the UCHIME program (47) referenced with the UNITE database (https://unite.ut.ee/) (48). Non-chimera sequences that passed the filtering processes were binned into operational taxonomic units (OTUs) at a 97% similarity cutoff using the UPARSE pipeline (47). All singletons OTU were excluded from the dataset. Subsequently, the taxonomic identity of each OTU was determined based on BLAST search against the UNITE reference database. The quality and depth of coverage of both primers reads were not significant difference, thus libraries from ITS1f reads were used for further analysis of fungal communities.

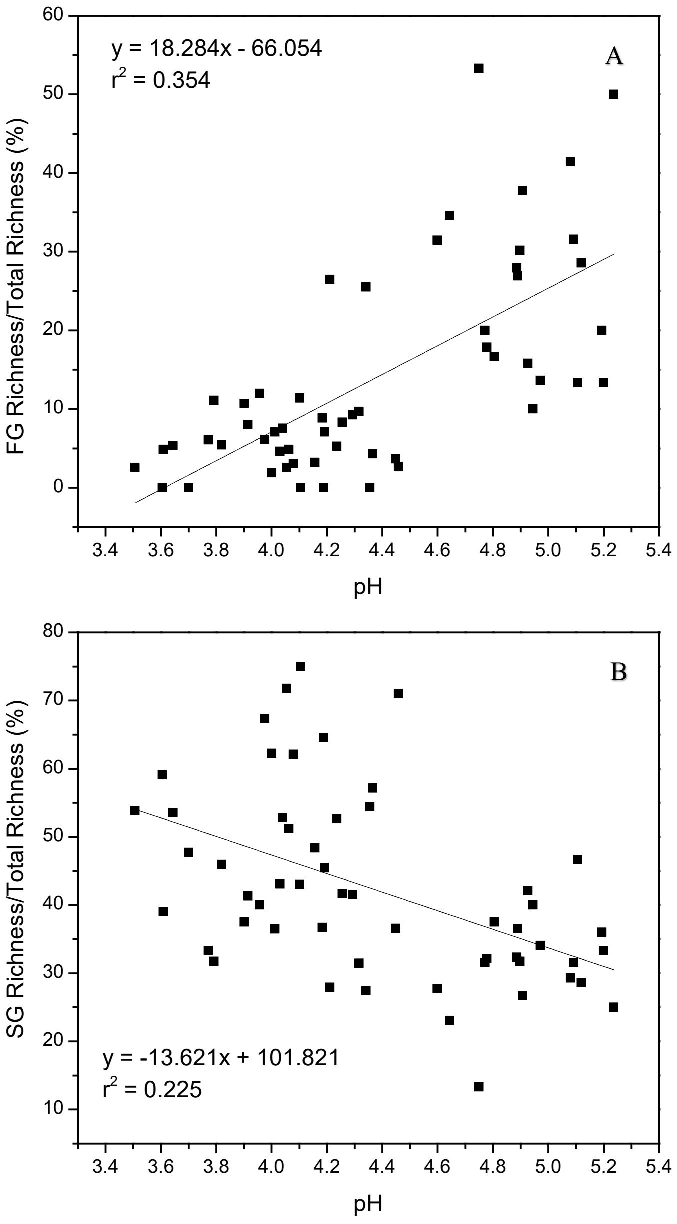


**Fig. S1** Effects of pH on ratios of (A) fast- and (B)slow-growing fungal richness to total richness. Species richness is the number of total OTUs observed in each sample. SG = slow-growing fungi, FG = fast-growing fungi

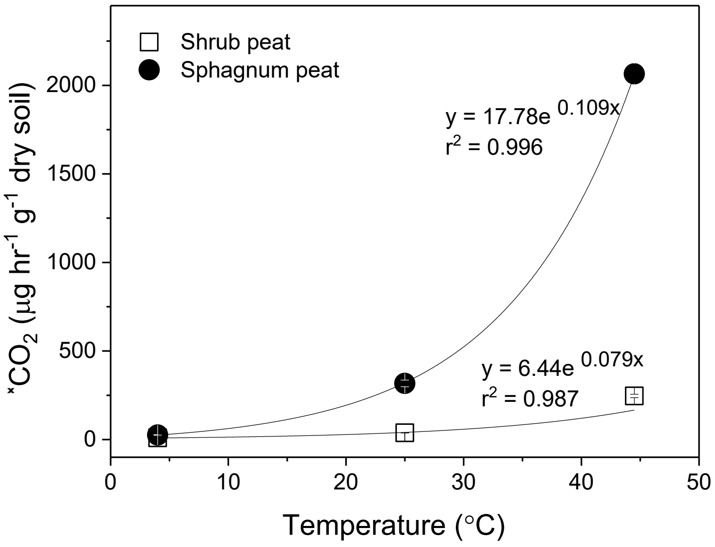


**Fig. S2** Temperature sensitivity of soil respiration (mean ± S.E.) in the subtropical shrub and the boreal *Sphagnum* peats through a lab incubation experiment. Regression analyses were run using $R = \alpha e^{\beta T}$, where R is soil respiration, coefficient α is the intercept of soil respiration when temperature is zero, coefficient β represents the temperature sensitivity of soil respiration and T is soil temperature

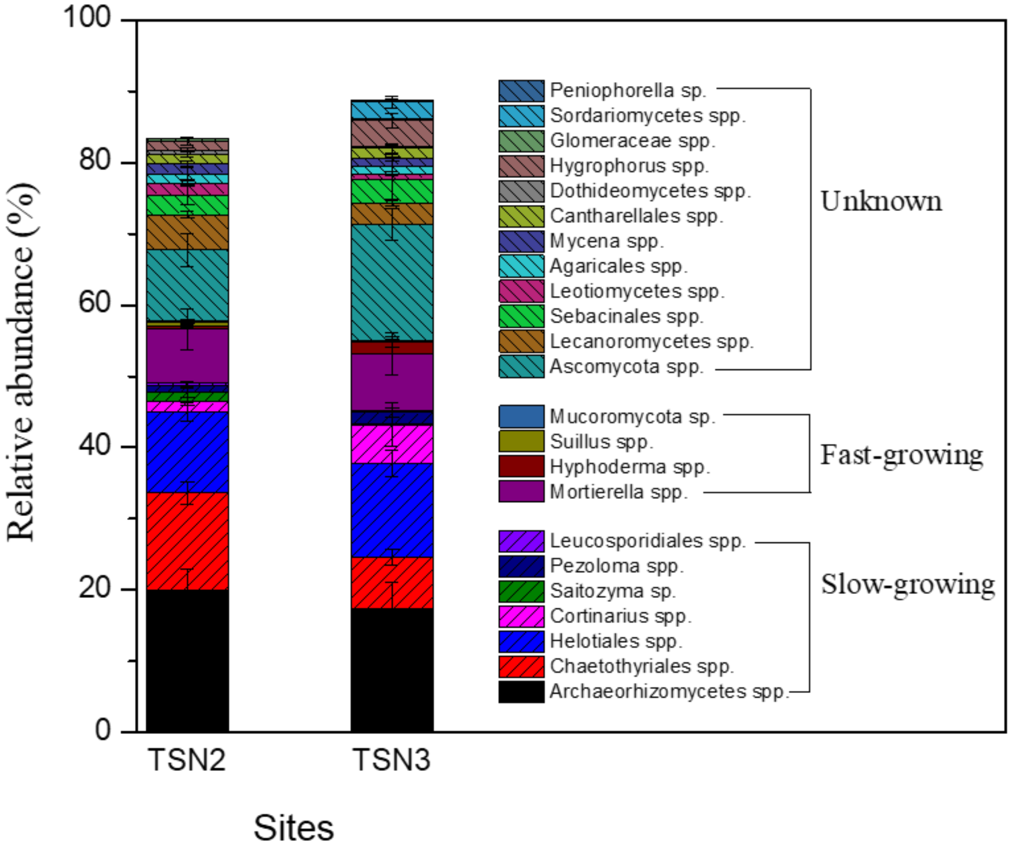


**Fig. S3** Relative abundance (mean ± S.E.) of the dominant fungi (OTU relative abundance >0.1%) in a bog forest (TSN2) and a *Sphagnum*-shrub mixed peat bog (TSN3) in the Pacific coastal temperate rainforest in Canada (36). The ecological traits of the dominant fungi present in Supplementary Table 1. The relative abundances were recalculated from raw amplicon reads in the European Nucleotide Archive [ITS(ERS1798771-ERS1799064)]

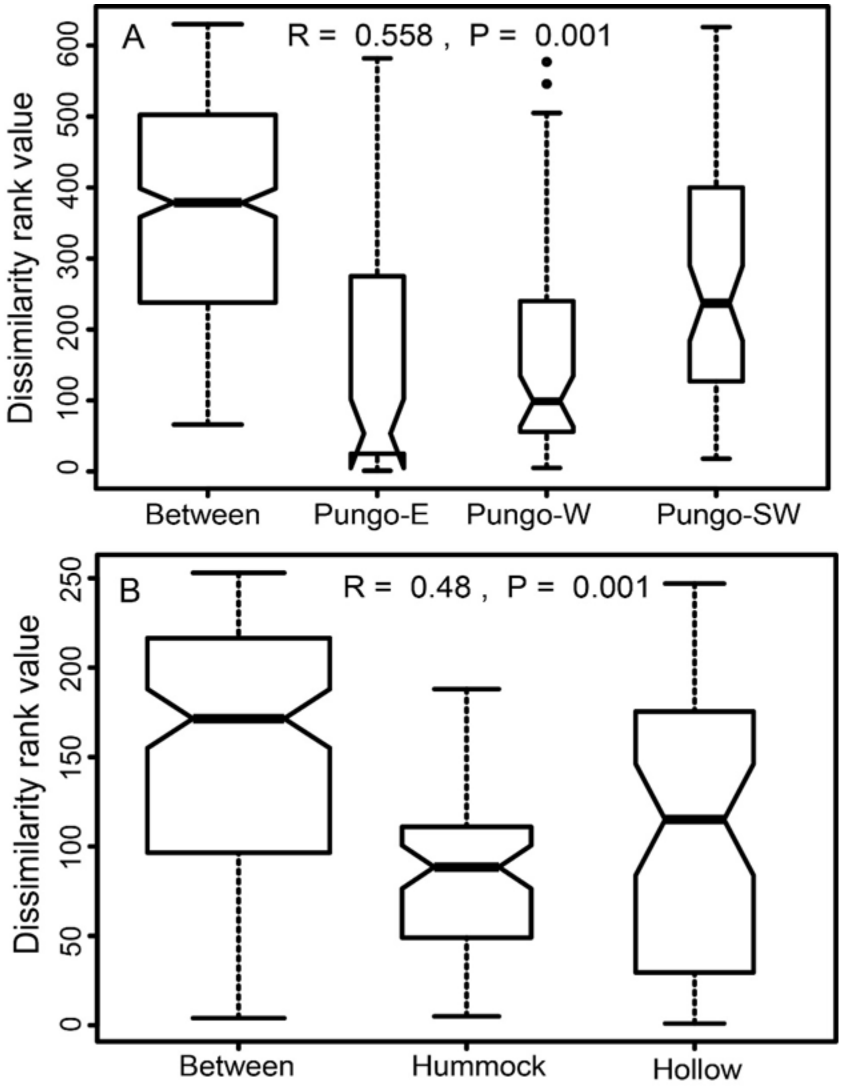


Fig. S4 Analysis of similarities (ANOSIM) plot showing dissimilarity between and within sites at (A) the shrub peatlands in NC and (B) the *Sphagnum* peatlands in MN. Bold horizontal bar in the box indicates median; bottom of the box indicates 25$^{th}$ percentile; top of the box indicates 75$^{th}$ percentile; whiskers extend to the most extreme data point, which is no more than the range (i.e. 1.5) times the interquartile range from the box; width of the bar is directly proportional to sample size.

Table S1 Characterization of research sites at Pocosin Lakes National Wildlife Refuge, Marcell Experimental Forest and Dajiuhu, add Canada sites

| | Pocosin Lakes National Wildlife Refuge, USA | | | Marcell Experimental Forest, USA | | Dajiuhu Peatlands,China | Calvert Island Field Station (ref. 36),Canada | |
|---|---|---|---|---|---|---|---|---|
| Sites | Pungo West | Pungo Southwest | Pungo-East | S1H4S1 (Hollow) | S1H4L1 (Hummock) | N/A | TSN2 (Bog forest) | TSN3 (*Sphagnum*/shrub bog) |
| Coordinate | 35.68882°, -76.56437° | 35.673478°, -76.5581° | 35.69078°, -76.52856° | 47.50871°, -93.45240° | 47.5084°, -93.45163° | 31.48333°, 109. 98333° | 51.65222°, -128.19272° | 51.65139°, -128.12861° |
| Dominant Species | *Ilex glabra, Lyonia lucida* | *Ilex glabra, Lyonia lucida, Woodwardia virginica* | *Lyonia lucida, Ilex glabra, Zenobia pulverulenta, Gaylussacia frondosa, Vaccinium formosum, Pinus serotina* | *S. fallax S. angustifolium, Picea mariana, Pinus resinosa, Larix laricina,* | *S. angustifolium, S. magellanicum, Ledum groenlandicum, Chamaedaphne calyculata, Vaccinium oxycoccos, Picea mariana, Pinus resinosa, Larix laricina,* | *Sphagnum palustre Crataegus wilsonii* | *Pinuscontorta, Chamaecyparis nootkatensis, andThuja plicata.`* | Abundant ericaceous shrubs and *Sphagnum* spp |
| Mean Water Level | *-39.6** | *-22.0** | *-30.5** | N/A | N/A | N/A | *-31.7* | *-40.7* |
| MAT(°C) | 16.8 | | | 3.3 | | 7.2 | N/A | N/A |
| Soluble Phenolics in soil (µg C/g dry soil) | 858±64 | 1171±98 | 658±137 | 114±11 | 107±21 | 89±26 | N/A | N/A |
| TIN (µg N/g dry soil) | 76±8 | 37±10 | 49±4 | 202±28 | 130±23 | 90±16 | 30.1±4.5 | 55.7±8.2 |
| TN | 1.89±0.08 | 1.88±0.07 | 1.11±0.04 | 1.42±0.14 | 1.35±0.24 | N/A | 1.13±0.04 | 1.40±0.05 |
| TC (%) | 55.91±0.78 | 51.33±0.71 | 50.09±0.93 | 43.69±1.02 | 44.52±2.58 | 38.0±7.4 | 55.5±0.9 | 57.6±1.1 |
| pH | 4.06±0.05 | 4.19±0.04 | 3.86±0.08 | 5.00±0.06 | 4.76±0.09 | 4.7±0.12 | 3.76±0.04 | 3.72±0.06 |
| SM (%) | 241±33 | 322±15 | 253±29 | 1214±91 | 1328±117 | 398±104 | `966±43 | 820±54 |

MAT = Mean Annual Temperature
TIN =Total Soil Inorganic Nitrogen ($NO_x$-N + $NH_4$-N)
TN =Total Soil Nitrogen
SM =Soil Moisture
TC = Total Soil Carbon
N/A = not available
* mean water level from 3/17/2015 to 10/05/2017 at Pocosin Lakes National Wildlife Refuge

Table S2 Growth rates of dominant fungi in the subtropical shrub and the boreal *Sphagnum* peatlands

| Division | Lower taxa | Growth rates | Dominant Fungi | | Ref. |
|---|---|---|---|---|---|
| | | | Shrub sites | *Sphagnum* site | |
| Ascomycota | *Archaeorhizomyces* spp. | Slow | + | − | (30, 49) |
| Ascomycota | *Pseudogymnoascus roseus* | Slow | − | + | (50) |
| Ascomycota | Helotiales sp. | Slow | + | − | (31, 51) |
| Basidiomycota | *Cryptococcus* sp.(= *Saitozyma* sp.) | Slow | + | − | (52-54) |
| Basidiomycota | *Solicoccozyma* sp. | Slow | + | − | (52) |
| Basidiomycota | *Guehomyces* sp. | Slow | + | − | (54) |
| Basidiomycota | *Cystofilobasidium* sp. | Slow | + | − | (52-54) |
| Basidiomycota | *Leucosporidiella* sp. | Slow | + | − | (53) |
| Ascomycota | *Calycina* sp. | Slow | + | − | (55) |
| Ascomycota | *Chaetothyriales* spp. | Slow | + | − | (56, 57) |
| Basidiomycota | *Cortinarius* spp | Slow | + | − | (58) |
| Ascomycota | *Pezoloma* spp. | Slow | + | − | (59) |
| Ascomycota | *Pseudeurotium* sp. | Fast | − | + | (60) |
| Ascomycota | *Penicillium* sp. | Fast | − | + | (61, 62) |
| Ascomycota | Saccharomycetales sp. | Fast | − | + | (63) |
| Basidiomycota | *Rhodotorula* sp. | Fast | − | + | (64) |
| Zygomycota | *Mortierella* spp. | Fast | − | + | (65, 66) |
| Basidiomycota | *Bullera* sp. | Fast | + | − | (67, 68) |
| Ascomycota | *Aspergillus* sp. | Fast | + | − | (29) |
| Basidiomycota | *Hyphoderma* spp. | Fast | + | − | (69) |
| Basidiomycota | *Suillus* spp. | Fast | + | − | (70) |
| Zygomycota | Mucoromycota sp. | Fast | + | − | (71) |
