## Supplementary Data 2 for "Vegetation and Microbes Interact to Preserve Organic Matter in Wooded Peatlands"

**Dataset 2 Mean percent of total reads from ITS4 primer for abundance >0.1% species or OTU (operational taxonomic unit, >0.1%) identified. The relative abundance > 1% were indicated in Bold.**

| Taxonomy/OTU | Pocosin Shrub peatland | | Minnesota *Sphagnum* peatland | |
|---|---|---|---|---|
| | Average | SD | Average | SD |
| *Pseudeurotium* sp | 0 | 0 | **0.447798** | **0.270642** |
| Archaeorhizomyces sp | **0.021743** | **0.012882** | 3.11E-05 | 6.21E-05 |
| *Oidiodendron maius* | 0.001893 | 0.002115 | 0 | 0 |
| Herpotrichiellaceae sp | 0.001029 | 0.001344 | 1.85E-05 | 3.7E-05 |
| Archaeorhizomyces sp | **0.029896** | **0.016006** | 7.19E-05 | 0.000125 |
| Inocybaceae sp | 0.001081 | 0.001178 | 0 | 0 |
| *Penicillium lividum* | 8.7E-06 | 1.74E-05 | 0.003196 | 0.003937 |
| Helotiaceae sp | 0 | 0 | 0.002625 | 0.003444 |
| Archaeorhizomyces sp | **0.019038** | **0.015012** | 0.000141 | 0.000108 |
| Archaeorhizomyces borealis | 0.001001 | 0.001857 | 0 | 0 |
| *Oidiodendron* sp | 3.46E-05 | 6.91E-05 | 0.001938 | 0.002529 |
| Archaeorhizomyces sp | **0.0374** | **0.022153** | 0.000157 | 0.000177 |
| Helotiales sp | **0.063455** | **0.016296** | 0.000559 | 0.000442 |
| Helotiales sp | 0.009021 | 0.01532 | 0 | 0 |
| Helotiales sp | 0.006984 | 0.004493 | 0 | 0 |
| Helotiales sp | 0.005485 | 0.003792 | 0 | 0 |
| Helotiales sp | 0.003047 | 0.006056 | 0 | 0 |
| Helotiales sp | 0.002263 | 0.003161 | 0 | 0 |
| Helotiales sp | 0.002308 | 0.002598 | 0 | 0 |
| Helotiales sp | 0.003698 | 0.007163 | 0 | 0 |
| *Mortierella* sp | 0 | 0 | **0.045498** | **0.024804** |
| *Entyloma microsporum* | 0 | 0 | 0.001986 | 0.002375 |

Notes: for all database, please contact Hongjun Wang,
